# BBSdb, an open resource for bacterial biofilm-associated proteins

**DOI:** 10.1101/2023.09.10.556831

**Authors:** Zhiyuan Zhang, Yuanyuan Pan, Wajid Hussain, Qingqing Li, Erguang Li

## Abstract

**Summary:** Bacterial biofilms are organized heterogeneous assemblages of microbial cells encased within a self-produced matrix of exopolysaccharides, extracellular DNA and proteins. Over the last decade, more and more biofilm-associated proteins have been discovered and investigated. Furthermore, omics techniques such as transcriptomes, proteomes also play important roles in identifying new biofilm-associated genes or proteins. However, those important data have been uploaded separately to various databases, which creates obstacles for biofilm researchers to have a comprehensive access to these data. In this work, we constructed BBSdb, a state-of-the-art open resource of bacterial biofilm-associated protein. It includes 48 different bacteria species, 105 transcriptome datasets, 21 proteome datasets, 1205 experimental samples, 57,823 differentially expressed genes (DEGs), 13,605 differentially expressed proteins (DEPs), 1,930 ‘Top 5% differentially expressed genes’ and a predictor for prediction of biofilm-associated protein. In addition, 1,781 biofilm-associated proteins, including annotation and sequences, were extracted from 942 articles and public databases via text-mining analysis. We used *E. coli* as an example to represent how to explore potential biofilm-associated proteins in bacteria. We believe that this study will be of broad interest to researchers in field of bacteria, especially biofilms, which are involved in bacterial growth, pathogenicity, and drug resistance.

**Availability and implementation:** The BBSdb is freely available at http://124.222.145.44/#!/.

## 1. Introduction

Bacterial biofilms are adhesion structure formed by single or multiple bacteria and their metabolites. In clinical practice, biofilms can greatly improve the ability of pathogenic bacteria to resist antibiotics, thus increasing the risk of infection (Schwarzer et al., 2020). Biofilm-associated proteins are defined as a type of protein molecules closely related to bacterial biofilm formation, which include constitutive proteins located downstream of the biofilm regulatory network and upstream transcriptional regulators. The understanding and discovery of biofilm-associated genes and proteins can help us to better understand the molecular mechanisms of bacterial biofilm formation. Over the last decade, more and more biofilm-associated genes and proteins have been discovered and investigated with the development of omics techniques including transcriptomes and proteomes (Lasaro et al., 2009; Hay et al., 2015; Wang et al., 2017; Jia et al., 2022). During the development and formation of biofilm, the transcription profile of bacteria changes, and some genes with obviously variable expression levels, which are proved by previous experiments, often play an important role in biofilm formation. Therefore, it is important for researchers to obtain these data and analyze gene and protein expression profile in the background of biofilms. However, those important transcriptome and proteome data have been uploaded separately to various databases, which make biofilm researchers pain to have a comprehensive access to these data. Although several resources provide biofilm data, such as Quorumpeps (Wynendaele et al., 2013) for QS-derived signaling peptides, BiofOmics (Lourenço et al., 2012) for biofilm experimental information, BaAMPs (Di et al., 2015), aBiofilm (Rajput et al., 2018), dpABB (Sharma et al., 2016) for antibiofilm Agents, BSD8 (Magalhães, et al., 2020) for structural information. There is an urgent need to combine multi-omics data for the prediction and analysis of biofilm-associated proteins.

Here, we developed BBSdb, an online database focusing on experimentally validated biofilm-associated proteins. In addition, BBSdb provided a predictor for prediction of biofilm-associated protein, in which users could upload their interested protein sequence to predict candidate biofilm-associated proteins and browse corresponding entries of DEG. BBSdb can serve as a useful resource to make researchers pain-free to obtain transcriptomes, proteomes in biofilm research, query information of experimentally validated biofilm-associated proteins, and utilize developed predictor for protein prediction.

## 2. Materials and methods

### 2.1 Data collection, pre-processing and analysis

To give users a clear study design of the data collection, pre-processing, analysis and integration in BBSdb, the overview of study design is illustrated in **Figure 1**.

**Figure 1.**
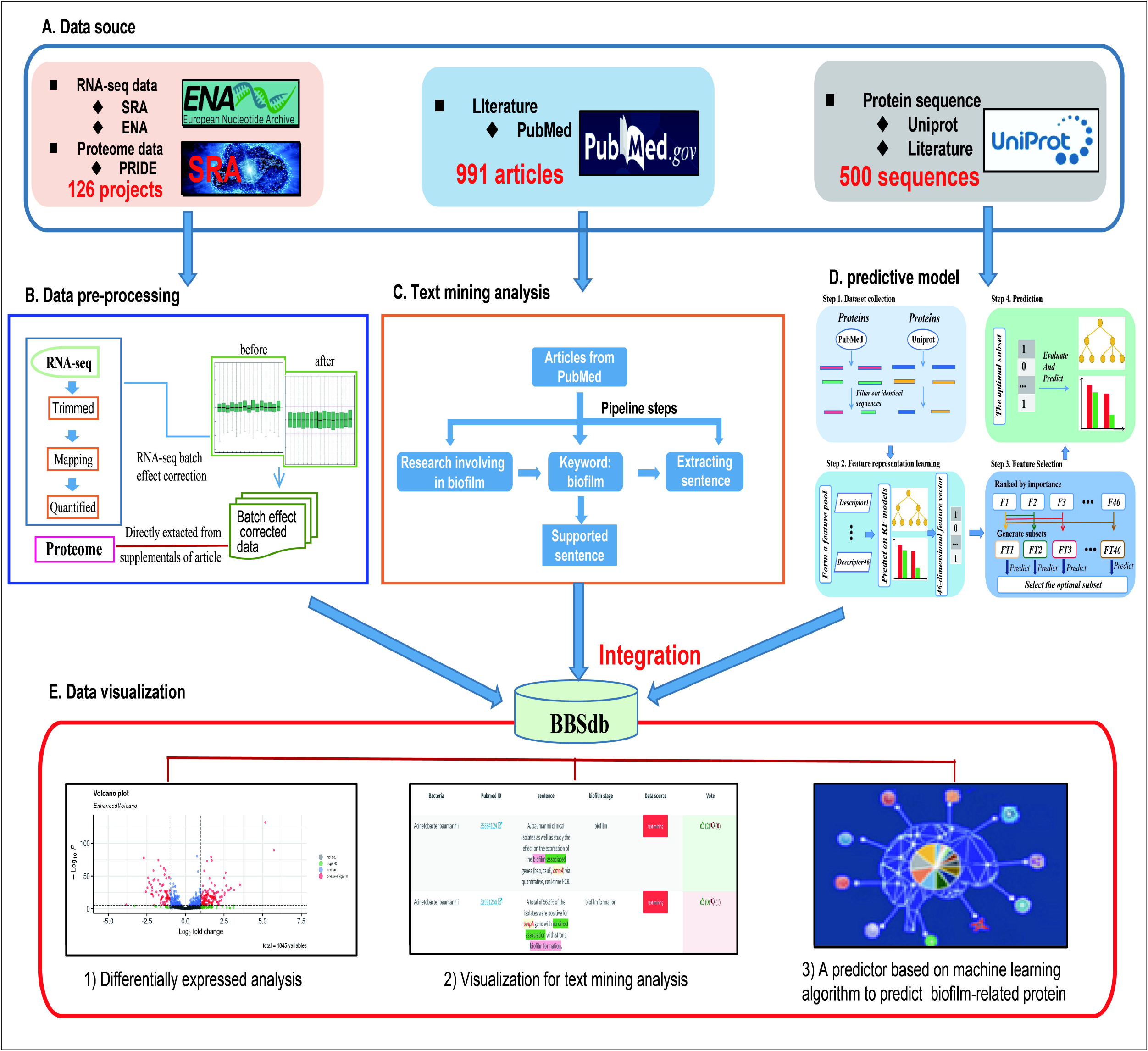
Overview of BBSdb. (A) The data source. All data including the raw sequencing data, information of genome annotation, literature, and experimental validated biofilm-associated proteins were collected from public databases, such as NCBI GEO, ENA, Uniprot, PubMed and so on. **(B)** The data pre-processing. The RNA-seq raw data perform quality control, mapping, transcript quantification and normalization to obtain high-quality data for further analysis. **(C)** Text mining analysis. A customized Python pipeline was used by searching for biofilm-associated proteins (genes) in the titles or abstracts of literature from the NCBI PubMed database. **(D)** Construction of predictive model. Firstly, the targeted protein sequences are subjected to the feature representation learning scheme, and as a result, a 46-dimensional feature vector will be generated. Secondly, the feature vector generated at the previous step is optimized to a 3-dimensional optimal feature vector. Ultimately, the proteins are predicted and scored by the well-trained RF model. **(E)** The data integration. BBSdb integrated polytype data with a biofilm-associated proteins predictor, which placed in “Bacteria”, “Genes(Proteins)”, “Diff. Expr.” and “Prediction” modules.

For data collection, we collected and curated transcriptome datasets from biofilm research, including 105 RNA-seq projects (1205 samples) from 16 different bacteria species, from the European Nucleotide Archive (EBI ENA, https://www.ebi.ac.uk/ena/browser/home) (Amid et al., 2020) and Sequence Read Archive (SRA, https://www.ncbi.nlm.nih.gov/sra/) (Kodama et al., 2012). As the related meta-data of corresponding experiments, projects and literature were obtained from NCBI PubMed and GEO databases (Barrett et al., 2013). For the processing of raw sequencing reads, FastQC (http://www.bioinformatics.babraham.ac.uk/projects/fastqc/) was used to evaluate the overall quality of the raw sequencing reads, followed by the Trim_galore to remove sequencing vectors and low-quality bases (Utturkar et al., 2020) and performed transcript quantification using Salmon (Patro et al., 2017), which adopted TPM (Transcript Per Million) for normalization (**Figure 1B**), a better unit for RNA abundance than RPKM and FPKM since it respects the invariance property and is proportional to the average relative RNA molar concentration (Zhao et al., 2020). For proteome datasets, we directly collected them from supplementary data of articles. We analyzed cleaned transcriptome and proteome data using differential expressed analysis, and obtaining 57,385 DEGs and 13,605 DEPs, in which we used a cutoff of |log2 FC| > 1.5 (FC, fold change) and p-value <0.05 to define differentially expressed genes and proteins between experiments (Zhang et al., 2019). In addition, we calculated ‘Top 5% differentially expressed gene’ of 16 different bacteria species, which was proposed by our previous research (Zhang et al., 2022).

### 2.2 Text-mining analysis

To obtain experimentally validated biofilm-associated proteins, we performed text-mining analysis. The main steps included: 1) obtaining literature, which must focus on bacterial biofilm at the gene or protein level; 2) obtaining sentences as supported evidence, which must describe gene or protein and contain the following keyword: biofilm; 3) extracting information of potential biofilm-associated gene or protein. The retrieved information was further verified by three rounds of manual inspection (**Figure 1C**). We obtained 1,781 experimentally validated biofilm-associated proteins, which were validated by 2,514 entries of supported evidence from 942 articles and public databases. We selected above-mentioned protein sequences with sequence similarity < 50% (CD-HIT) as positive datasets for following predictor development. And reviewed non biofilm-associated proteins from Uniprot database (https://www.uniprot.org/) (UniProt, Consortium, 2021) were extracted as negative datasets (http://124.222.145.44/#!/terms).

### 2.3 Predictor development

The work used a feature representation learning scheme that integrated different types of sequence-based feature descriptors to develop predictive model (**Figure 2A**). For protein sequence-based feature descriptors, we utilized different feature extraction methods (Rajput et al., 2015; Wei et al., 2020) for feature representation and obtained 46 feature descriptors in total (Supplementary S1). We compared all 46 feature descriptors using RF (Random Forest) classifier, and the top 3 best feature descriptors CKSAAP (Tung et al., 2013), TPC (Duan et al., 2008), and AAC_DPC with the RF classifier obtained 0.7 0.72, 0.75 AUC scores, respectively (**Figure 2B**). We constructed predictive model based on a 3-dimensional feature vector, which consisted of prediction result of CKSAAP, TPC, and AAC_DPC descriptors on datasets, and obtained an AUC score 0.80 (**Figure 2C-D**). BBSdb provided ‘Prediction’ module to make users free for biofilm-associated protein prediction.

**Figure 2.**
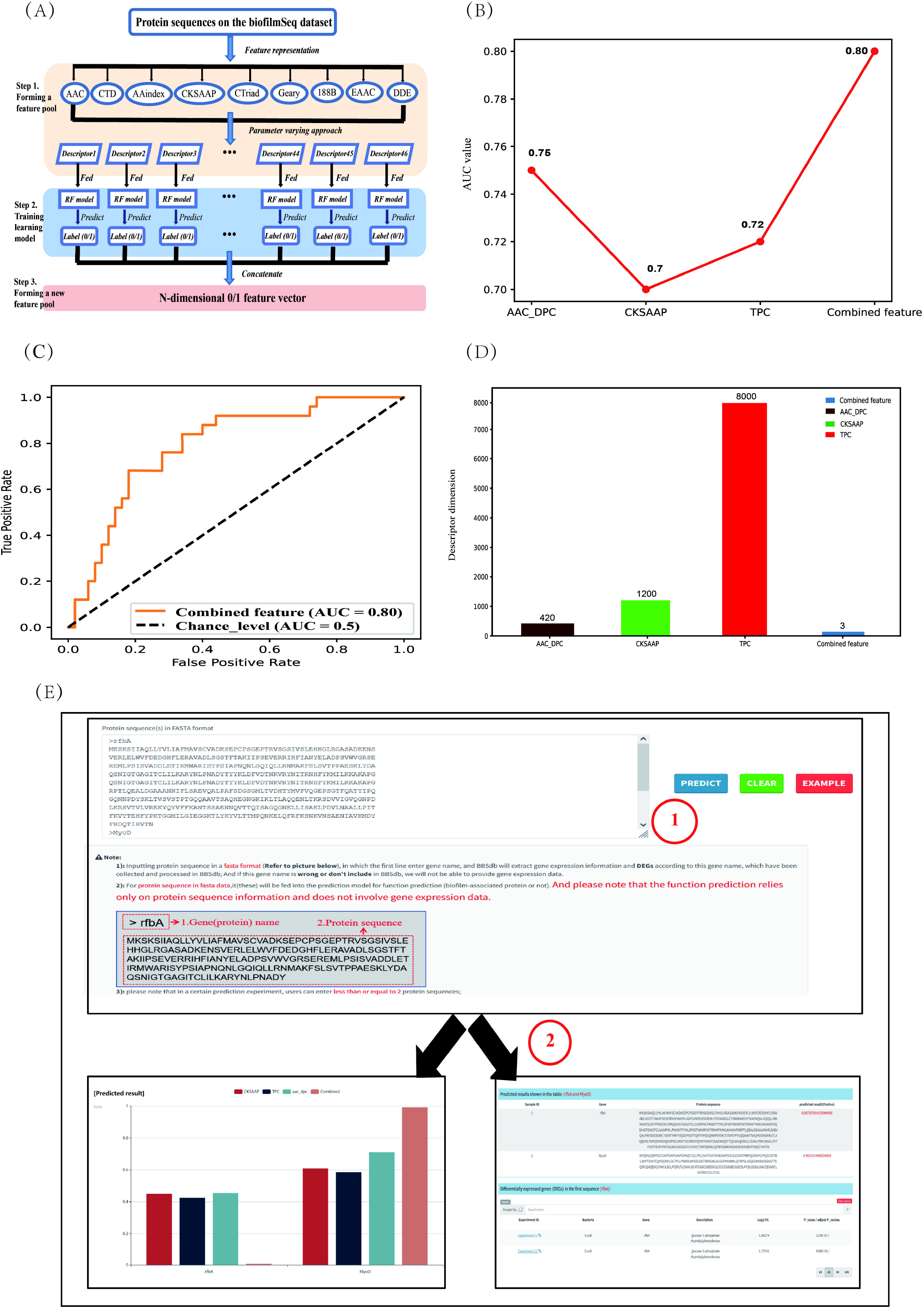
Introduction of predictor for biofilm-associated protein. (A) Pipeline of the feature representation learning scheme. Firstly, a feature pool with 46 feature descriptors is constructed by nine feature-encoding algorithms. Afterwards, each descriptor is trained and evaluated using the RF classifier on the biofilm-associated protein sequence datasets. Finally, the predicted class label for each trained RF model is regarded as an attribute to form a new feature vector. **(B)** Comparative analysis of single feature descriptor with combined feature (feature subsets). **(C)** ROC curves of the best-performing feature descriptors using RF classifier. **(D)** Feature dimension of the optimal feature (combined feature) and The original three feature descriptors. **(E)** The introduction of “Prediction” module. Users can enter their protein sequences of interest (in fasta format) into the input box on the module page, and then click the ‘Predict’ button on the right. At this point, the entered protein sequence is fed into the prediction model for function prediction, while the protein name is used to search for corresponding gene expression information in BBSdb database.

### 2.4 Database design and implementation

BBSdb, which integrated data and a predictor, was designed as a relational database. All data were loaded into a MySQL database. The frontend of the website was coded using JavaScript and HTML, while the backend was coded using Python with a Flask framework to support queries to the MySQL database and provide representational state transfer (REST) application programming interfaces (APIs) for programmable access to our data. The AngularJS framework was used to bride the frontand back-ends. Echarts.js and plotly.js used for visualizations at the front end.

## 3. Results

### 3.1 Summary of BBSdb

The BBSdb database provides a user-friendly, open access web interface for searching, browsing and downloading data, which includes 105 transcriptome projects from 16 different bacteria species and 21 proteome projects from 5 different bacteria species and 1,781 entries of experimentally validated biofilm-associated protein via text-mining analysis, which were validated by 2,514 entries of supported evidence from 942 articles and public databases. We analyzed RNA-seq and proteome datasets and performed differentially expressed analysis, in total obtaining 33,180 DEGs, 13,605 DEPs. We calculated ‘Top 5% differentially expressed genes’ of 16 different bacteria species, and obtaining 1,930 ‘Top 5% differentially expressed genes’. All data shown in **Table 1**.

**Table 1.**
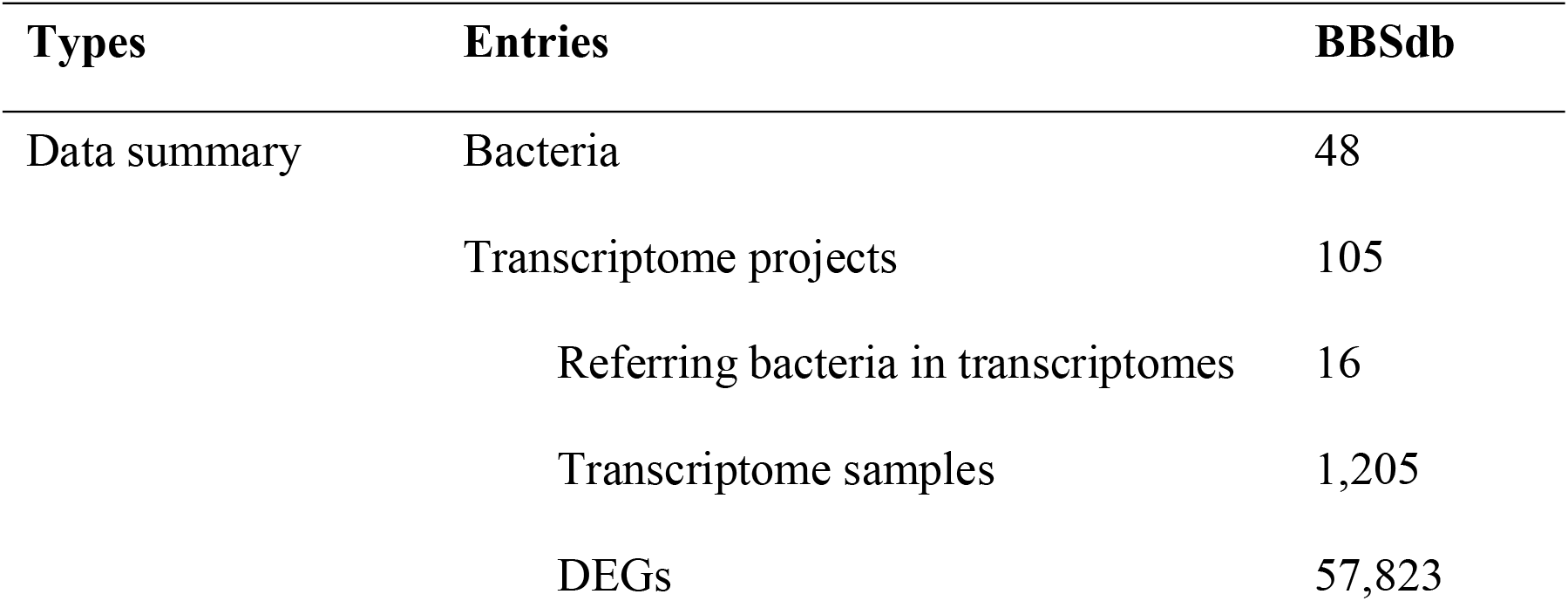
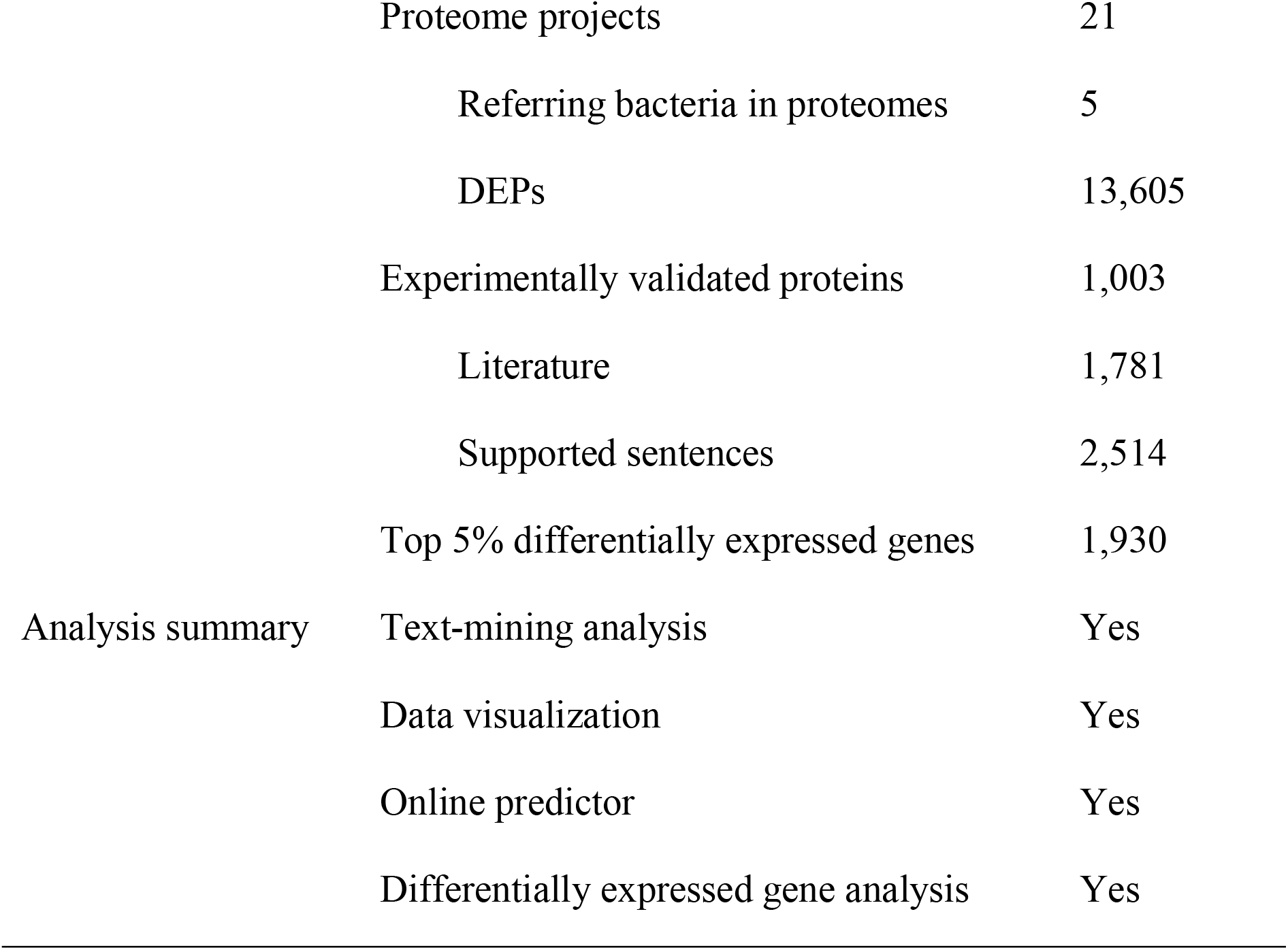
Summary in BBSdb database.

### 3.2 Effective online predictor

‘Prediction’ module aims to provide an online predictor for identification of potential biofilm-associated protein. Users can input protein sequence in a fasta format, in which the first line enter gene name, and BBSdb will extract gene expression information and DEGs according to inputted gene name, which have been collected and processed in BBSdb (**Figure 2E**); And if this gene name is wrong or don’t include in BBSdb, gene expression data will not be able to be provided. For protein sequence of the second line, it will be fed into the prediction model for function prediction (biofilm-associated protein or not). And please note that the function prediction relies only on protein sequence information and does not involve in gene expression data.

### 3.3 Functional modules

The BBSdb database mainly includes four functional modules, ‘Bacteria’, ‘Genes (Proteins)’, ‘Diff. Expr.’, and ‘Proteome’. The ‘Bacteria’ module consists of two tables (**Figure 3A, 3E**), with the first table displaying biofilm-associated protein information from 48 bacteria species, as reported in the literature or recorded in public databases. The second table summarizes transcriptomic information from 16 bacteria species, including the number of experimental samples, the number of DEGs, and links providing more detailed information. Users can obtain transcriptomic experimental information of bacteria, DEGs, and experimentally validated biofilm-associated proteins by clicking on the link. The ‘Genes (Proteins)’ module aims to provide researchers with experimentally validated biofilm-associated proteins and corresponding evidence obtained through data-mining analysis, allowing users to select the bacteria and genes of interest to retrieve information (**Figure 3B, 3F)**. In addition, we provided a user-feedback mechanism, allowing users to vote on each text mining entry. Users can use the thumb-up and thumb-down buttons to vote on the creditability of the entry, the background color of the table cell will be changed automatically according to the voting results. In ‘Diff. Expr.’ module, we provided DEGs and specific information on transcriptomic experiments and visualized these data to facilitate users’ querying and mining (**Figure 3C, 3G**). The ‘Proteome’ module collected proteomic data from 5 bacteria species and provided the analyzed differentially expressed proteins. We placed the results of the combined analysis of bacterial transcriptomics and proteomics in this module, and provided their overlapping genes or proteins (**Figure 3D**), which help us to find new regulatory factors under the bacterial biofilm phenotype through data mining.

**Figure 3.**
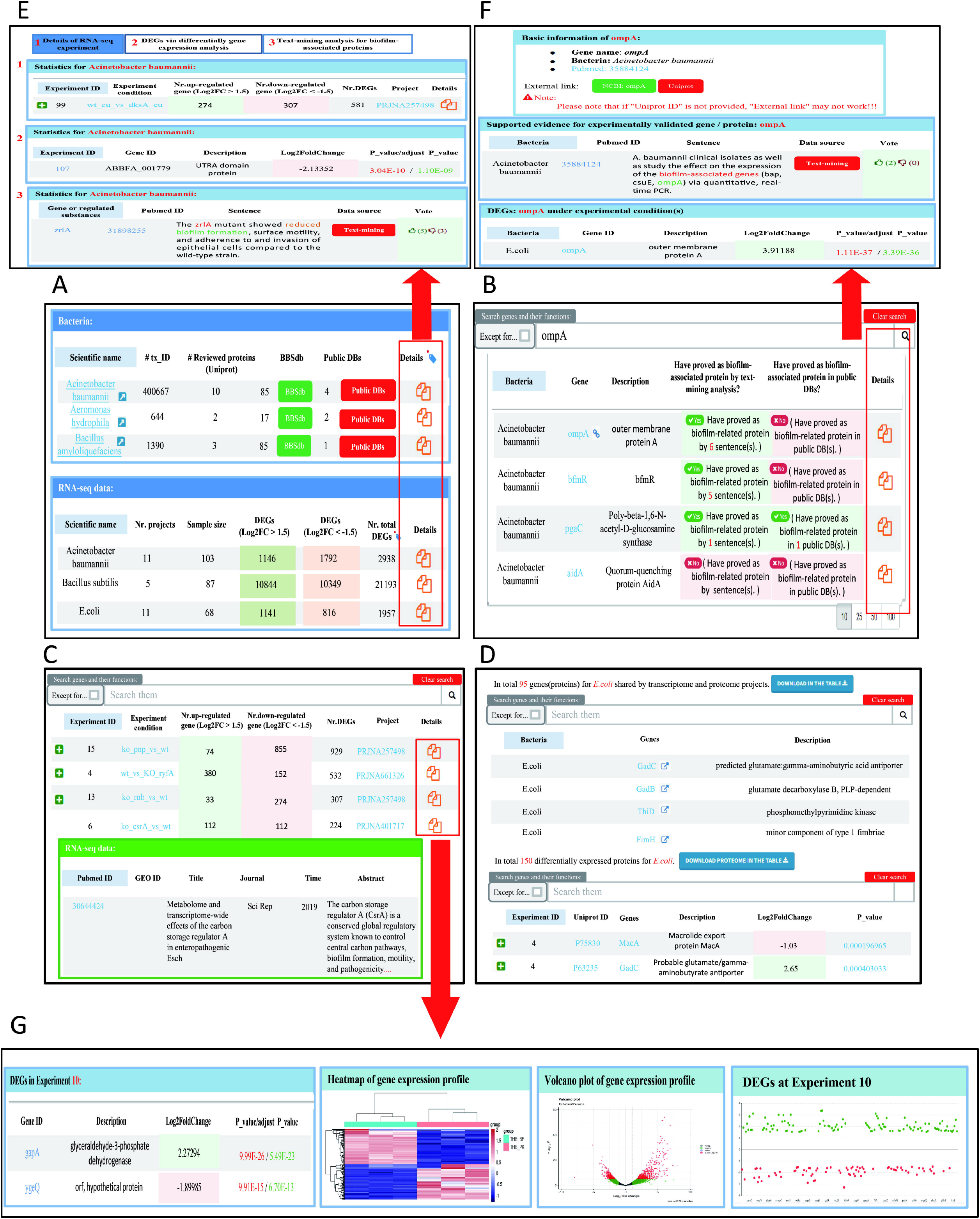
Introduction of four main functional module of BBSdb database. (A) ‘Bacteria’ module of BBSdb database. This module summarized experimentally validated biofilm-associated protein information from 48 different bacteria species and transcriptomic information from 16 different bacteria species. **(B)** ‘Genes (Proteins)’ module of BBSdb database. This module provided researchers with experimentally validated biofilm-associated proteins and evidence obtained through data-mining analysis. (**C)** ‘Diff. Expr.’ module of BBSdb database. This module provided transcriptomic data and DEGs from 16 different bacteria species, and visualized these data to facilitate users’ querying and mining of DEGs. (**D)** ‘Proteome’ module of BBSdb database. This module provided proteomic data and DEPs from 5 different bacteria species. In addition, BBSdb performed multi-omics combined analysis, obtained overlapping genes and proteins between transcriptomics and proteomics were integrated in this module. **(E)** The bacteria detail page. Users can obtain detailed transcriptomic experimental information of bacteria, DEGs, and experimentally validated biofilm-related proteins. **(F)** The Genes (Proteins) detail page. **(G)** The Diff. Expr. detail page.

### 3.4 Multi-omics data mining

We classified 16 bacteria species into two groups: (1) Human microbial pathogens, in total 15: *E*.*coli, Burkholderia cenocepacia, Streptococcus pneumoniae, Streptococcus mutans, Staphylococcus epidermidis, Staphylococcus aureus, Salmonella enterica, Pseudomonas aeruginosa, Porphyromonas gingivalis, Listeria monocytogenes, Klebsiella pneumoniae, Acinetobacter baumannii, Mycobacterium Tuberculosis, Vibrio cholerae*, and *Listeria monocytogenes*; (2) others: *Bacillus subtilis*. For human pathogenic bacteria, we analyzed the gene expression of different bacteria species, and found that in the background of biofilm, the number of DEGs of different bacteria species are quite different, which may be due to the number of genes in the bacterial genome itself is different. The proportion of DEGs to the number of genes in the bacterial genome range from 0.3% to 53% (**Figure 4A**). We also analyzed gene expression in single type of bacteria under different conditions. According to differential expression analysis, the shared genes that were differentially expressed under different experimental conditions and ‘Top 5% differentially expressed genes’ were obtained, which might play important role in biofilm phenotype. Due to there being only one transcriptome dataset for *Vibrio cholerae* and *Pseudomonas aeruginosa*, their ‘Top 5% differentially expressed genes’ could not be obtained. For the 14 different bacteria species *Klebsiella pneumoniae, E*.*coli, Bacillus subtilis, Acinetobacter baumannii, Enterococcus faecalis, Staphylococcus aureus, Porphyromonas gingivalis, Burkholderia cenocepacia, Staphylococcus epidermidis, Streptococcus mutans, Streptococcus pneumoniae, Salmonella enterica*, the number of ‘Top 5% differentially expressed genes’ are 297, 283, 230, 221, 190, 171, 152, 148, 73, 73, 52, 28, 12, separately (**Figure 4B**), which are likely involved in the regulatory development of the biofilm as genes encoding biofilm-associated proteins. Subsequently, we performed a multi-omics analysis based on transcriptomic and proteomic data, and identified 95, 418, 367, 1958 shared proteins (genes) in *E*.*coli, Staphylococcus aureus, Pseudomonas aeruginosa*, and *Bacillus subtilis*, respectively (**Figure 4C**); By enriching these shared genes (proteins) in the ‘Top 5% differentially expressed genes’ datasets of the bacteria, we obtained 26, 47, 34, 128 entries respectively (**Figure 4D**); STRING (Szklarczyk et al., 2023), GO (Gene Ontology, Consortium, 2015), and KEGG (Kanehisa et al., 2017) analysis revealed that for *Pseudomonas aeruginosa*, the 34 genes are enriched in biofilm formation and the type VI secretion system pathways (**Figure 4E**), suggesting their involvement in the regulatory development of biofilms, providing researchers with clues for a more comprehensive understanding of the biofilm developmental mechanism.

**Figure 4.**
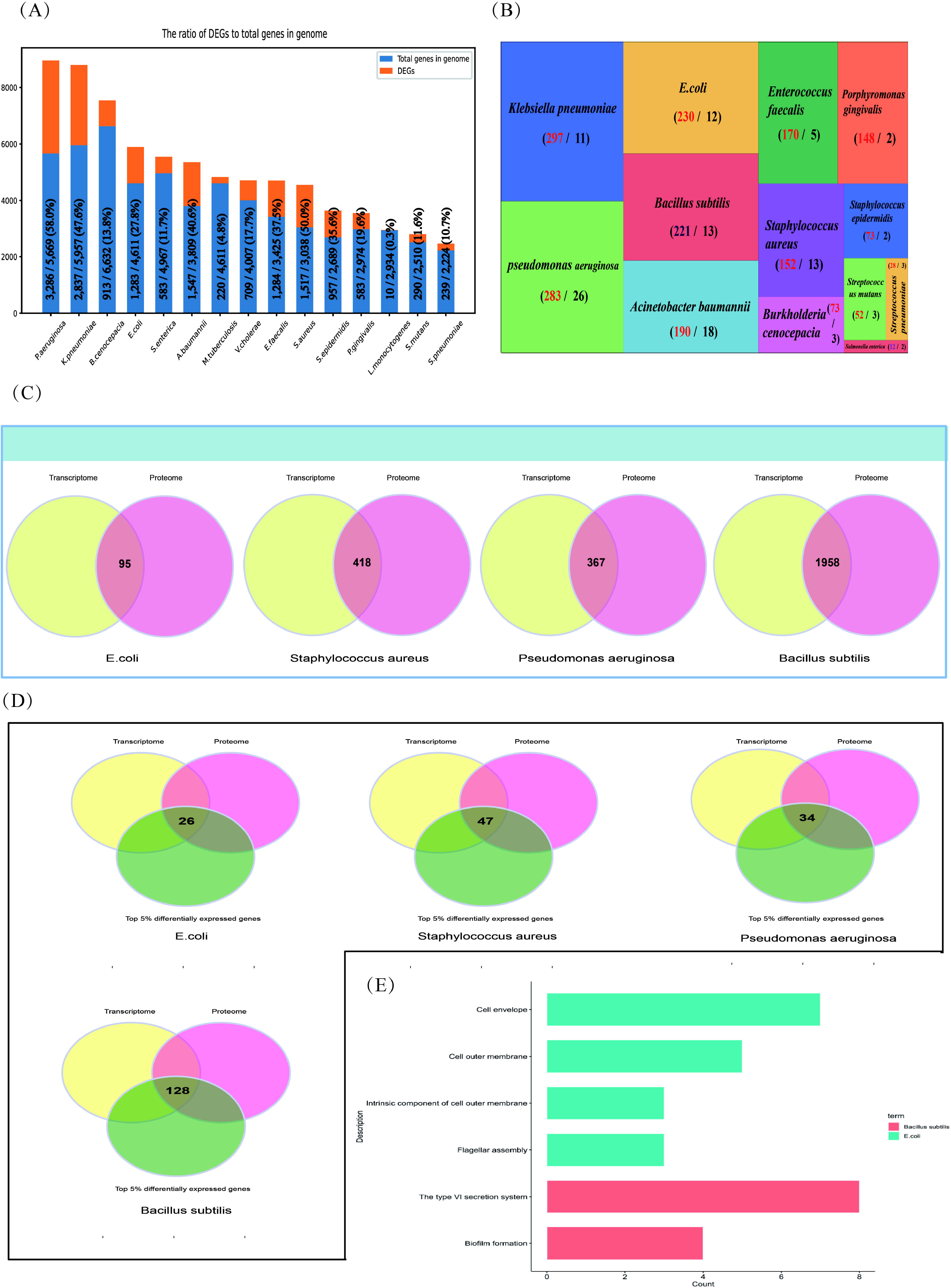
Multi-omics Data mining. **(A)** The number of DEGs of 15 Human microbial pathogens, and the proportion of DEGs to the number of genes in the bacterial genome. **(B)** Summary of Top 5% differentially expressed genes from 14 different bacteria species. BBSdb calculates the numbers of the experimental condition of each gene when differentially expressed in experiments, and ranks the corresponding gene by the calculated number of experimental conditions. **(C)** A multi-omics combined analysis between transcriptomics and proteomics to obtained shared genes (proteins) from 4 different bacteria species, including *E*.*coli, Staphylococcus aureus, Pseudomonas aeruginosa* and *Bacillus subtilis*. **(D)** The overlapping analysis between above-mentioned shared genes and ‘Top 5% differentially expressed genes’. **(E)** STRING, GO, KEGG analysis.

### 3.5 Case study

Users could use BBSdb to obtain existing biofilm-associated protein information and also to systematically explore potential biofilm-associated proteins utilizing collected data and developed tool. We used *E. coli* as an example to represent how to screen out biofilm-associated proteins using transcriptome data and developed predictor. Firstly, we calculated DEGs of 3 RNA-seq experiments for *E. coli* (**Experiment_1**: aerobic biofilm cultures versus aerobic planktonic cultures, **Experiment_2**: anaerobic biofilm cultures versus anaerobic planktonic cultures, and **Experiment_3**: biofilm versus planktonic cultures), in which the expression abundances were normalized as TPM values, using a cutoff of |log2 FC| > 1.5 (FC, fold change) and p-value <0.05 to define DEGs between experiments. We found that the number of DEGs in Experiment_1, Experiment_2, and Experiment_3 were 604, 1475, and 2217, respectively. Among these experiments, the number of shared genes of Experiment_1 with Experiment_2, Experiment_2 with Experiment_3, and Experiment_1 with Experiment_3 were 188, 1329, and 279, respectively (**Figure 5A**). In total, 168 shared genes were obtained from the 3 RNA-seq experiments. Meanwhile, we constructed protein-protein interaction (PPI) networks for 168 shared genes using STRING (**Figure 5B**), a current collection of known and predicted direct physical binding and indirect functionally related interactions between proteins / genes. The results indicated that these genes were involved in propanoate, butanoate and pyruvate metabolism. When exploring the top expression genes for the three experiments, we found that they shared 23 genes in the top 500 DEGs, including one commonly down-regulated gene *garP* (**Figure 5C**), which was associated with the plasma membrane, and 22 commonly upregulated genes (*b1551, infA, tyrU, cspI, groS, csrA, exbD, argQ, greA, sraF, metU, b4140, yifE, rplU, yheL, secG, zapA, wzzB, ydfK, cspB, cspG*, and *yjcB*) in three experiments (**Figure 5D**), which are mainly were involved in RNA-binding activity. Subsequently, we filtered out 4 tRNA genes (*tyrU, sraF, metU, argQ*) and then used the constructed classifier to predict 19 remaining DEGs, in total, 10 genes (*garP, yjcB, yifE, yheL, secG, b1551, csrA, zapA, ydfK, b4140*) were predicted to be biofilm-associated proteins (**Figure 1E**). For gene *zapA*, we noted that it is described in the literature as “Biofilm production was significantly associated with the expression of *zapA* (P < 0.05)”, which indicated *zapA* may be involved in biofilm formation, although its experimental subject was *Proteus mirabilis* (Sun et al., 2020). SecG belongs to the accessory Sec system (Bandara et al., 2016), and deleting member *secA2* of the Sec system caused a substantial reduction in biofilm formation. According to research, Asp5, which is necessary for early stage biofilm formation in *Streptococcus gordonii*, is homologous to SecG which indicated a similar SecG function (Bandara et al., 2016; Takamatsu et al., 2005). In addition, gene *csrA* plays an important role in biofilm formation by *E. coli* (Jackson et al., 2002) and *yifE* is involved in biofilm formation (https://www.uniprot.org/uniprotkb/P0ADN2/entry).

**Figure 5.**
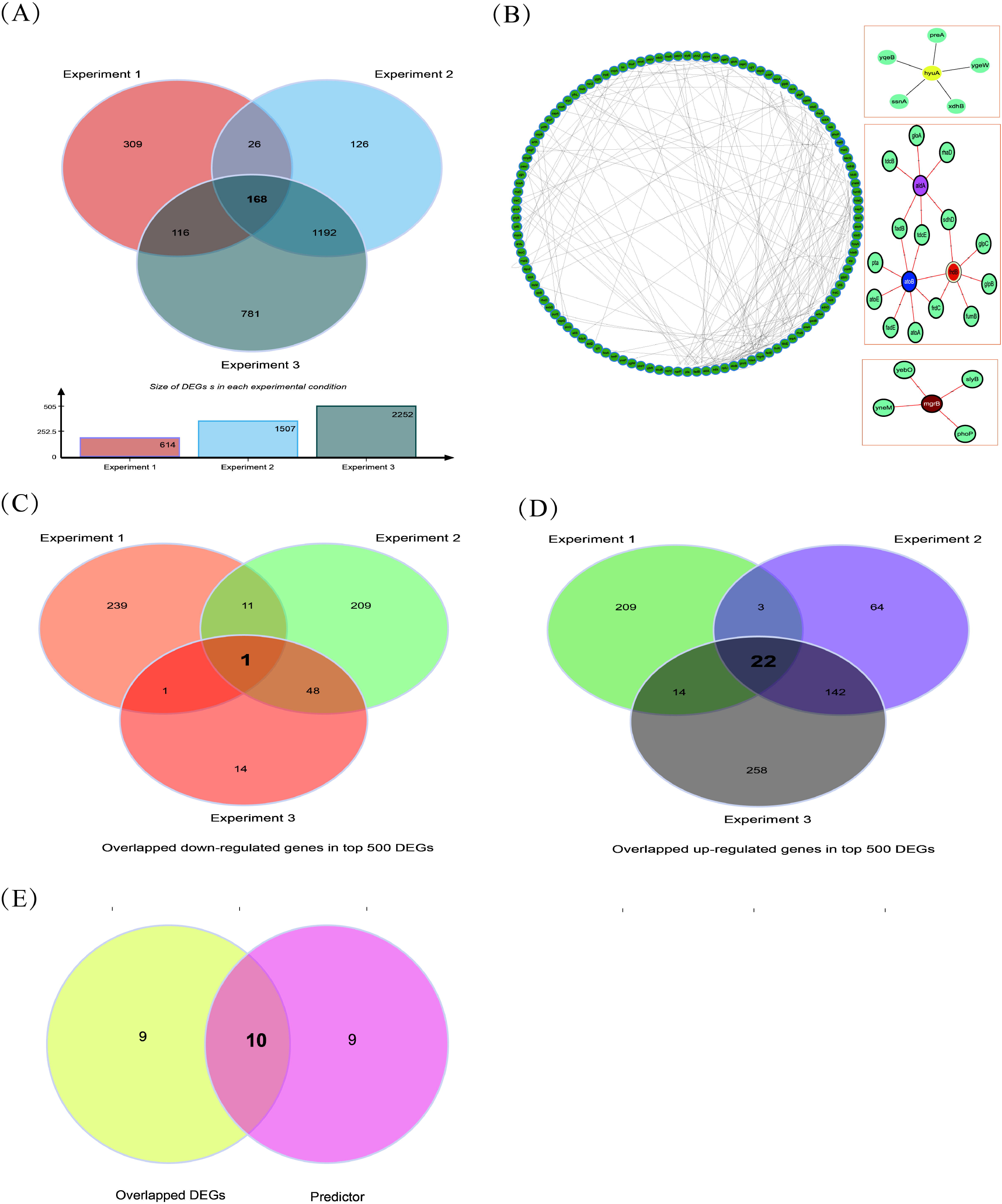
Case study. **(A)** The number of DEGs in Experiments1, 2, and 3 of 3 RNA-seq experiments for *E. coli* were 604, 1475, and 2217, respectively **(B)** The shared and specific DEGs of 3 RNA-seq experiments for *E. coli*. **(C)** The analysis of protein-protein interaction (PPI) interaction networks for 173 shared genes obtained from 3 RNA-seq experiments. **(D)** Top 500 DEGs were obtained by differentially expressed analysis for each RNA-seq experiment, then, up-regulated genes were retained to screen out overlapped up-regulated genes of three experiments. **(E)** Same as above, the down-regulated genes were retained to screen out overlapped down-regulated genes of three experiments.

## 4. Discussion

In this work, we provided a comprehensive database for biofilm research, which could be an infrastructure for the biofilm research community. In addition, we developed a predictor assisting researchers to explore potential biofilm-associated proteins. However, several questions remain to be addressed. For example, the BBSdb database only collected bacterial proteomes through Supplementary materials in literature without collecting, processing, and analyzing the original bacterial proteome data in public databases such as PRIDE (Perez-Riverol et al., 2022). Regarding the existence of multiple post-transcriptional regulation, proteomes will provide useful information for researchers, therefore more proteomes should also be integrated in the BBSdb.

Biofilm-associated proteins are participating in the biofilm formation process of their respective bacteria referring to previous research (Lasa et al., 2006). In this study, we defined biofilm-associated protein as those that have been reported in the literature and validated experimentally as components of the biofilm structure or involved in the regulation of biofilm development. In practical applications, we need to take into account the differences between different biofilm model systems and many other factors, such as the duration of the experiment and the specific strain, in order to obtain condition-dependent biofilm-associated proteins (Edel et al., 2019; Flemming et al., 2023). These proteins can effectively explain the diversity of mechanisms in bacterial biofilm development. In addition, homologous genes and proteins of different bacteria species can perform different functions, it is necessary to develop a bacteria-specific predictive model for better guiding practice in the next release.

In the future, other questions also remain to be addressed, including the long-term maintenance and update of BBSdb and improvement of predictive ability upon big data accumulation.

## Code availability

The source code for the developed platform is available at the BBSdb GitHub repository (https://github.com/895548727/Bacterial_biofilm).

