## Supplementals for "BBSdb, an open resource for bacterial biofilm-associated proteins"

**Open** **resource of bacterial biofilm for the prediction of biofilm-related proteins**

**Supplementary Material**

**Content**

#### 1. Used Feature Descriptors for Protein sequences

#### 1.1 Amino Acid Composition (AAC)

To include the compositional information, we computed the frequencies and the occurrences of 20 different amino acids in the peptide sequence, respectively. The feature vector for the AAC descriptor is represented below:

AAC(k)=(V1,V2,⋯,Va,…,V20),

where Va denotes the value of the amino acid type a, and k(k=1,2) denotes the method type; for example, 1 denotes the frequency while 2 denotes the occurrence number.

#### 1.2 Enhanced Amino Acid Composition (EAAC)

The Enhanced Amino Acid Composition (EAAC) feature type is introduced here for the first time. It calculates the AAC based on the sequence window of fixed length (the default value is 5) that continuously slides from the N- to C-terminus of each peptide and can be usually applied to encode the peptides with an equal length. An illustrated example of this encoding scheme is provided in the following **Figure S1**.

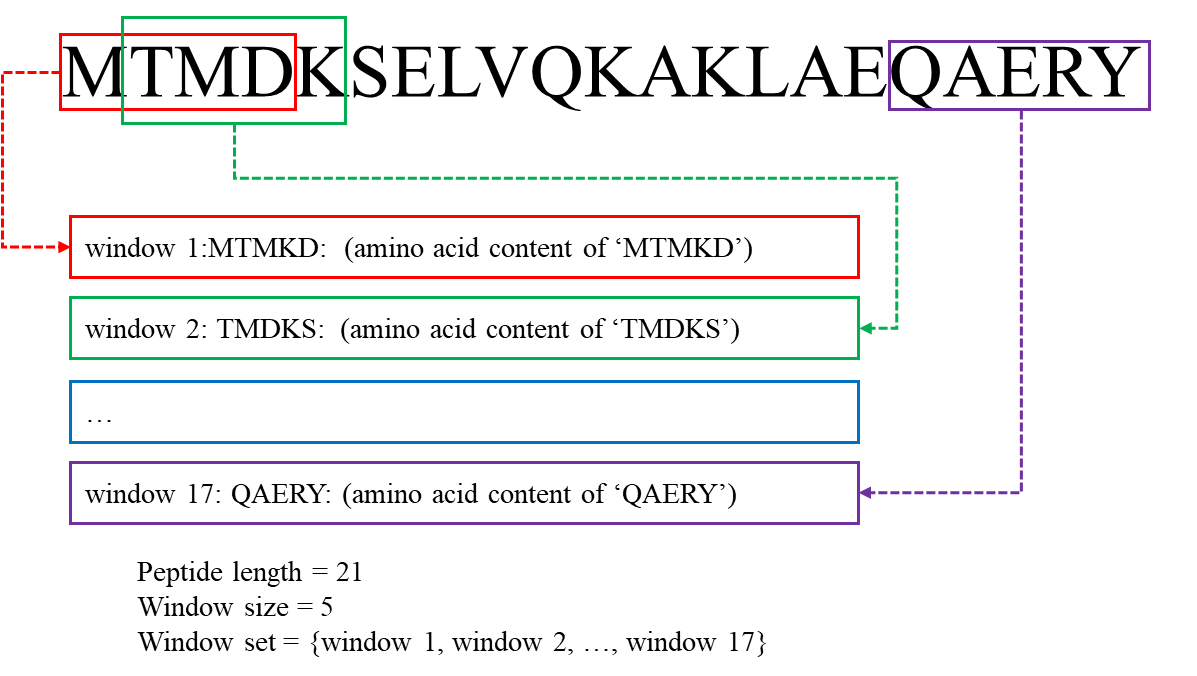

**Figure S1**. An illustrated example of the EAAC descriptor.

The EAAC can be calculated as:

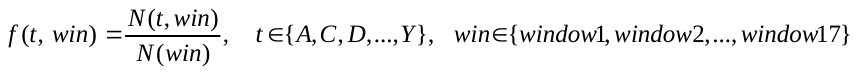

where *N(t,win)* is the number of amino acid type *t* in the sliding window *win* and *N(win)* is the size of the sliding window *win*.

### 1.3 Composition of *k*-spaced Amino Acid Pairs (CKSAAP)

The CKSAAP feature encoding calculates the frequency of amino acid pairs separated by any *k* residues (*k* = 0, 1, 2, … , 5. The default maximum value of *k* is 5) (Chen, et al., 2009; Chen, et al., 2007a; Chen, et al., 2007b; Chen, et al., 2008). Taking *k* = 0 as an example, there are 400 0-spaced residue pairs (i.e., AA, AC, AD,…, YY.). Then, a feature vector can be defined as:

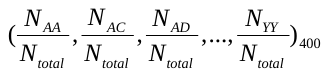

The value of each descriptor denotes the composition of the corresponding residue pair in the protein or peptide sequence. For instance, if the residue pair AA appears *m* times in the protein, the composition of the residue pair AA is equal to *m* divided by the total number of *0*-spaced residue pairs (*N_total_*) in the protein. For *k* = 0, 1, 2, 3, 4 and 5, the value of *N_total_* is *P* – 1, *P* – 2, *P* – 3, *P* – 4, *P* – 5 and *P* – 6 for a protein of length *P*, respectively. An illustrated example of this encoding scheme (*k*=0) is provided in the following **Figure S2**.

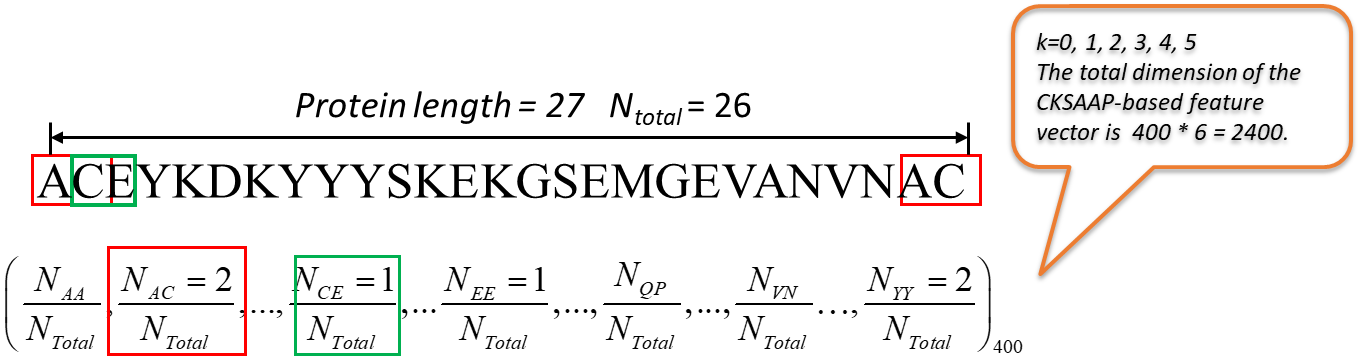

### 1.4 Tri-Peptide Composition (TPC)

The Tripeptide Composition (TPC) (Bhasin and Raghava, 2004) gives 8000 descriptors, defined as:

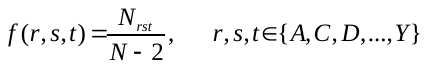

where *N_rst_* is the number of tripeptides represented by amino acid types *r*, *s* and *t.*

### 1.5 Di-Peptide Composition (DPC)

The Dipeptide Composition (Saravanan and Gautham, 2015) gives 400 descriptors. It is defined as:

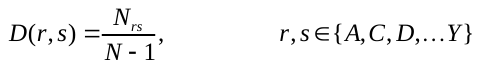

where *N_rs_* is the number of dipeptides represented by amino acid types *r* and *s.*

The Dipeptide Deviation from Expected Mean feature vector (Saravanan and Gautham, 2015) is constructed by computing three parameters, i.e. dipeptide composition (*D_c_*), theoretical mean (*T_m_*), and theoretical variance (*T_v_*). The above three parameters and the DDE are computed as follows. *D_c_(r,s)*, the dipeptide composition measure for the dipeptide ‘*rs’*, is given as

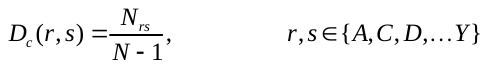

where *N_rs_* is the number of dipeptides represented by amino acid types *r* and *s* and *N* is the length of the protein or peptide. *T_m_(r,s)*, the theoretical mean, is given by:

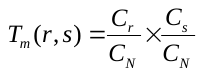

where *C_r_* is the number of codons that code for the first amino acid and *C_s_* is the number of codons that code for the second amino acid in the given dipeptide ‘*rs*’. *C_N_* is the total number of possible codons, excluding the three stop codons (i.e., 61). *T_v_* (*r,s*), the theoretical variance of the dipeptide ‘*rs*’, is given by:

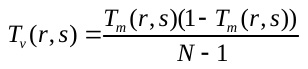

Finally, *DDE(r,s)* is calculated as:

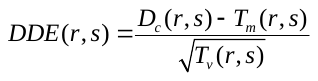

### 1.6 Grouped Amino Acid Composition (GAAC)

In the GAAC encoding, the 20 amino acid types are further categorized into five classes according to their physicochemical properties, e.g. hydrophobicity, charge and molecular size (Lee, et al., 2011b). The five classes include the aliphatic group (*g1*: GAVLMI), aromatic group (*g2*: FYW), positive charge group (*g3*: KRH), negative charged group (*g4*: DE) and uncharged group (*g5*: STCPNQ). GAAC descriptor is the frequency of each amino acid group, which is defined as:

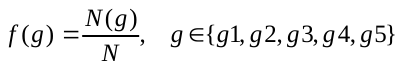

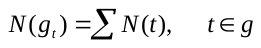

where *N(g)* is the number of amino acid in group *g*, *N(t)* is the number of amino acid type *t,* and *N* is the length of the protein/peptide sequence.

### 1.7 Enhanced GAAC (EGAAC)

The Enhanced GAAC (EGAAC) is also for the first time proposed in this work. It calculates GAAC in windows of fixed length (default is 5) continuously sliding from the N- to C-terminal of each peptide and is usually applied to peptides with an equal length.

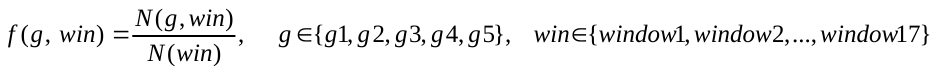

where *N(g, win)* is the number of amino acids in group *g* within the sliding window *win* and *N(win)* is the size of the sliding window *win*.

### 1.8 Composition of *k*-Spaced Amino Acid Group Pairs (CKSAAGP)

The Composition of *k*-Spaced Amino Acid Group Pairs (CKSAAGP) is a variation of the CKSAAP descriptor, which is our own proposal. It calculates the frequency of amino acid group pairs separated by any *k* residues (the default maximum value of *k* is set as 5). Taking *k* = 0 as an example, there are 25 0-spaced group pairs (i.e., *g1g1, g1g2, g1g3, … g5g5*). Thus, a feature vector of CKSAAGP can be defined as:

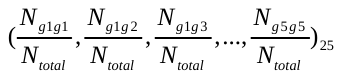

The value of each descriptor denotes the composition of the corresponding residue group pair in a protein or peptide sequence. For instance, if the residue group pair *g1g1* appears *m* times in the protein, the composition of the residue pair *g1g1* is equal to *m* divided by the total number of *0*-spaced residue pairs (*N_total_*) in the protein. For *k* = 0, 1, 2, 3, 4 and 5, the values of *N_total_* are *P* – 1, *P* – 2, *P* – 3, *P* – 4, *P* – 5 and *P* – 6 respectively, for a protein of length *P*.

### 1.9 Grouped Di-Peptide Composition (GDPC)

The Grouped Di-Peptide Composition encoding is another variation of the DPC descriptor. It is composed of a total of 25 descriptors that are defined as:

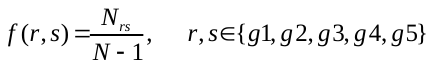

where *N_rs_* is the number of tripeptides represented by amino acid type groups *r* and *s*, *N* is the length of a protein or peptide sequence.

### 1.10 Grouped Tri-Peptide Composition (GTPC)

The Grouped Tri-Peptide Composition encoding is also a variation of TPC descriptor, which generates 125 descriptors, defined as:

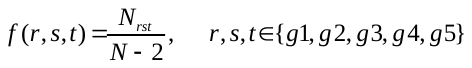

where *N_rst_* is the number of tripeptides represented by amino acid type groups *r*, *s* and *t*. *N* is the length of a protein or peptide sequence.

### 1.11 Geary correlation (Geary)

The Geary autocorrelation descriptors (Sokal and Thomson, 2006) for a protein or peptide sequence are defined as:

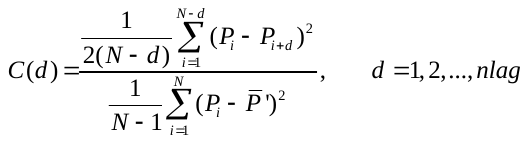

where *d*, *P*, *P_i_* and *P_i+d_*, *nlag* have the same definitions as described above.

### 1.12 Composition/Transition/Distribution (CTD)

The Composition, Transition and Distribution (CTD) features represent the amino acid distribution patterns of a specific structural or physicochemical property in a protein or peptide sequence (Cai, et al., 2003; Cai, et al., 2004; Dubchak, et al., 1995; Dubchak, et al., 1999; Han, et al., 2004). 13 types of physicochemical properties have been previously used for computing these features. These include hydrophobicity, normalized Van der Waals Volume, polarity, polarizability, charge, secondary structures and solvent accessibility. These descriptors are calculated according to the following procedures: (i) The sequence of amino acids is transformed into a sequence of certain structural or physicochemical properties of residues; (ii) Twenty amino acids are divided into three groups for each of the seven different physicochemical attributes based on the main clusters of the amino acid indices of Tomii and Kanehisa (Tomii and Kanehisa, 1996). The groups of amino acids are listed in **Table S1**.

**Table S1.** Amino acid physicochemical attributes and the division of the amino acids into three groups according to each attribute.

| Attribute | Division | | |
| --- | --- | --- | --- |
| Hydrophobicity_PRAM900101 | Polar: RKEDQN | Neutral: GASTPHY | Hydrophobicity: CLVIMFW |
| Hydrophobicity_ARGP820101 | Polar: QSTNGDE | Neutral: RAHCKMV | Hydrophobicity: LYPFIW |
| Hydrophobicity_ZIMJ680101 | Polar: QNGSWTDERA | Neutral: HMCKV | Hydrophobicity: LPFYI |
| Hydrophobicity_PONP930101 | Polar: KPDESNQT | Neutral: GRHA | Hydrophobicity: YMFWLCVI |
| Hydrophobicity_CASG920101 | Polar: KDEQPSRNTG | Neutral: AHYMLV | Hydrophobicity: FIWC |
| Hydrophobicity_ENGD860101 | Polar: RDKENQHYP | Neutral :SGTAW | Hydrophobicity: CVLIMF |
| Hydrophobicity_FASG890101 | Polar: KERSQD | Neutral: NTPG | Hydrophobicity: AYHWVMFLIC |
| Normalized van der Waals volume | Volume range: 0-2.78  GASTPD | Volume range: 2.95-94.0  NVEQIL | Volume range: 4.03-8.08  MHKFRYW |
| Polarity | Polarity value: 4.9-6.2  LIFWCMVY | Polarity value: 8.0-9.2  PATGS | Polarity value: 10.4-13.0  HQRKNED |
| Polarizability | Polarizability value: 0-1.08  GASDT | Polarizability value: 0.128-120.186  GPNVEQIL | Polarizability value: 0.219-0.409  KMHFRYW |
| Charge | Positive: KR | Neutral: ANCQGHILMFPSTWYV | Negative: DE |
| Secondary structure | Helix: EALMQKRH | Strand: VIYCWFT | Coil: GNPSD |
| Solvent accessibility | Buried: ALFCGIVW | Exposed: PKQEND | Intermediate: MPSTHY |

**1.12.1**	**CTDC**

Taking the hydrophobicity attribute as an example, all amino acids are divided into three groups: polar, neutral and hydrophobic (**Table S1**). The Composition descriptor consists of three values: the global compositions (percentage) of polar, neutral and hydrophobic residues of the protein. An illustrated example of this encoding scheme is provided in the following **Figure S4**. The Composition descriptor can be calculated as follows:

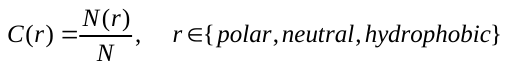

where *N(r)* is the number of amino acid type *r* in the encoded sequence and *N* is the length of the sequence.

### **1.12**.2 CTDT

The Transition descriptor T also consists of three values (Dubchak, et al., 1995; Dubchak, et al., 1999): A transition from the polar group to the neutral group is the percentage frequency with which a polar residue is followed by a neutral residue or a neutral residue by a polar residue. Transitions between the neutral group and the hydrophobic group and those between the hydrophobic group and the polar group are defined in a similar way. The transition descriptor can then be calculated as:

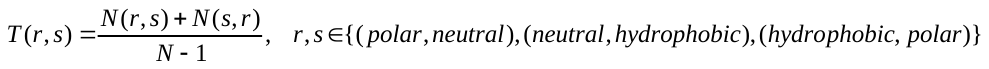

where *N(r,s)* and *N(s,r)* are the numbers of dipeptides encoded as “*rs*” and “*sr*” respectively in the sequence, while *N* is the length of the sequence. An illustrated example of this encoding scheme is provided in the following **Figure S4**.

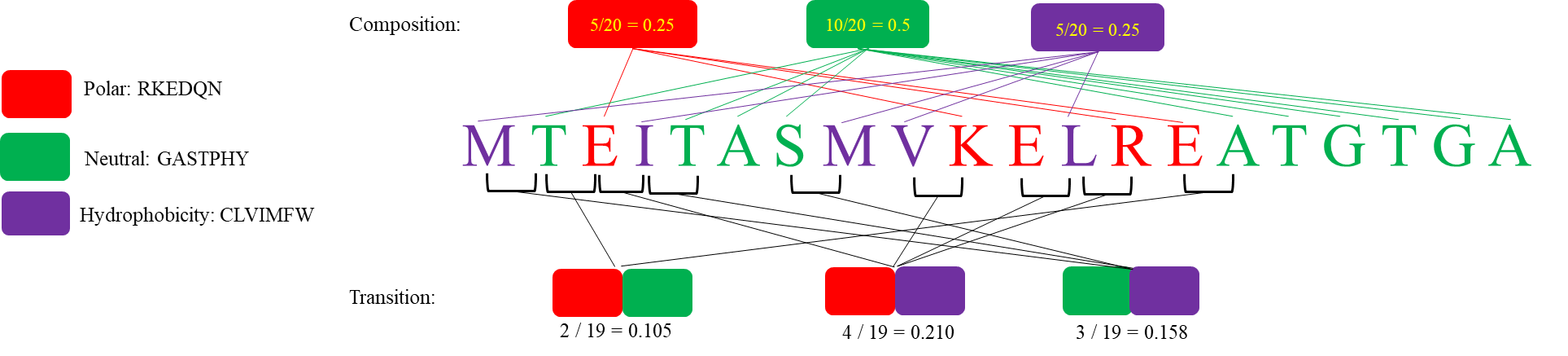

**Figure S4**. An illustrated example of the calculation of composition and transition descriptors. This example uses the hydrophobicity attribute.

### **1.12.**3 CTDD

The Distribution descriptor consists of five values for each of the three groups (polar, neutral and hydrophobic) (Dubchak, et al., 1995; Dubchak, et al., 1999), namely the corresponding fraction of the entire sequence, where the first residue of a given group is located, and where 25, 50, 75 and 100% of occurrences are contained.

For example, we start with the first residue up to and including the residue that marks 25/50/75/100% of occurrences for residues of any given group and then we simply divide the position of this residue by the length of the entire sequence.

### 1.13 Conjoint Triad (CTriad)

The Conjoint Triad descriptor (CTriad) considers the properties of one amino acid and its vicinal amino acids by regarding any three continuous amino acids as a single unit (Shen, et al., 2007). First, the protein sequence is represented by a binary space (*V, F*), where *V* denotes the vector space of the sequence features, and each feature (*V_i_*) represents a sort of triad type; *F* is the number vector corresponding to *V*, where *f_i_*, the value of the *i-*th dimension of *F*, is the number of type *V_i_* appearing in the protein sequence.

For the amino acids that have been catalogued into seven classes, the size of *V* should be equal to 7ⅹ7ⅹ7=343. Accordingly, *i* = 1, 2, 3, …, 343. An illustrated example of this encoding scheme is provided in the following **Figure S5**.

In principle, the longer a protein sequence, the higher the probability to have larger values of *f_i_*, confounding the comparison of proteins with different lengths. Thus, we define a new parameter, *d_i_*, by normalizing *f_i_* with the following equation:

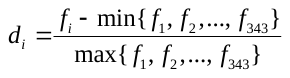

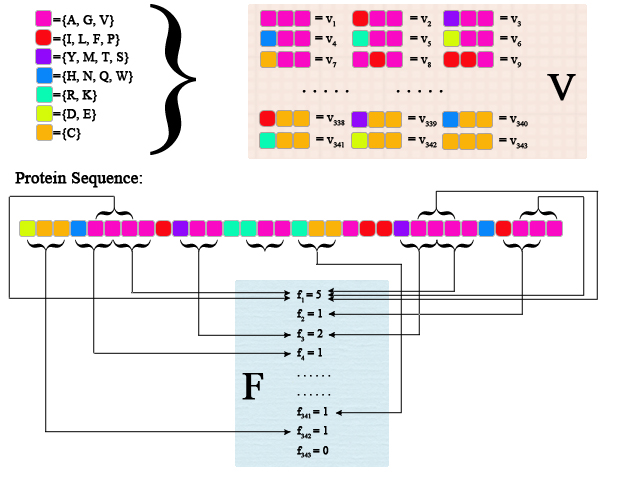

**Figure S5**. Schematic diagram for constructing the vector space (*V, F*) of a given protein sequence. *V* is the vector space of the sequence features; each feature (*V_i_*) represents a triad composed of three consecutive amino acids; *F* is the number vector corresponding to *V*, and the value of the *i-*th entry of *F*, denoted *f_i_*, is the number of occurrences that the triad associated with *V_i_* appearing in the protein sequence. The figure was adapted from the Supplementary Figure in (Shen, et al., 2007).

### 1.14 Pseudo-Amino Acid Composition (PAAC)

This group of descriptors has been proposed in (Chou, 2001; Chou, 2005). Let $H_{1}^{o}(i)$, $H_{2}^{o}(i)$,$M^{o}\left( i \right)$ for *i* = 1, 2, 3, … 20 be the original hydrophobicity values, the original hydrophilicity values and the original side chain masses of the 20 natural amino acids, respectively. They are converted to the following quantities by a standard conversion:

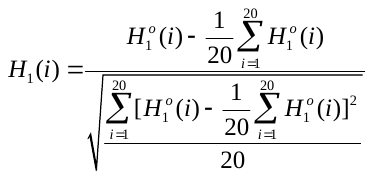

where $H_{2}^{o}(i)$ and $M^{o}(i)$ are normalized as $H_{2}(i)$ and $M(i)$ in the same manner.

Next, a correlation function can be defined as:

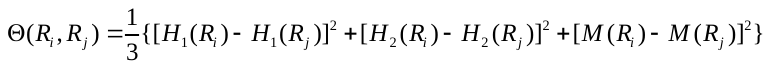

This correlation function is actually an averaged value for the three amino acid properties: hydrophobicity value, hydrophilicity value and side chain mass. Therefore, we can extend this definition of correlation function for one amino acid property or for a set of *n* amino acid properties.

For one amino acid property, the correlation can be defined as:

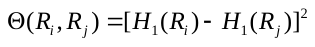

where *H(R_i_)* is the amino acid property of amino acid *R_i_* after standardization.

An illustrated example of the correlation function is provided in the following **Figure S8**.

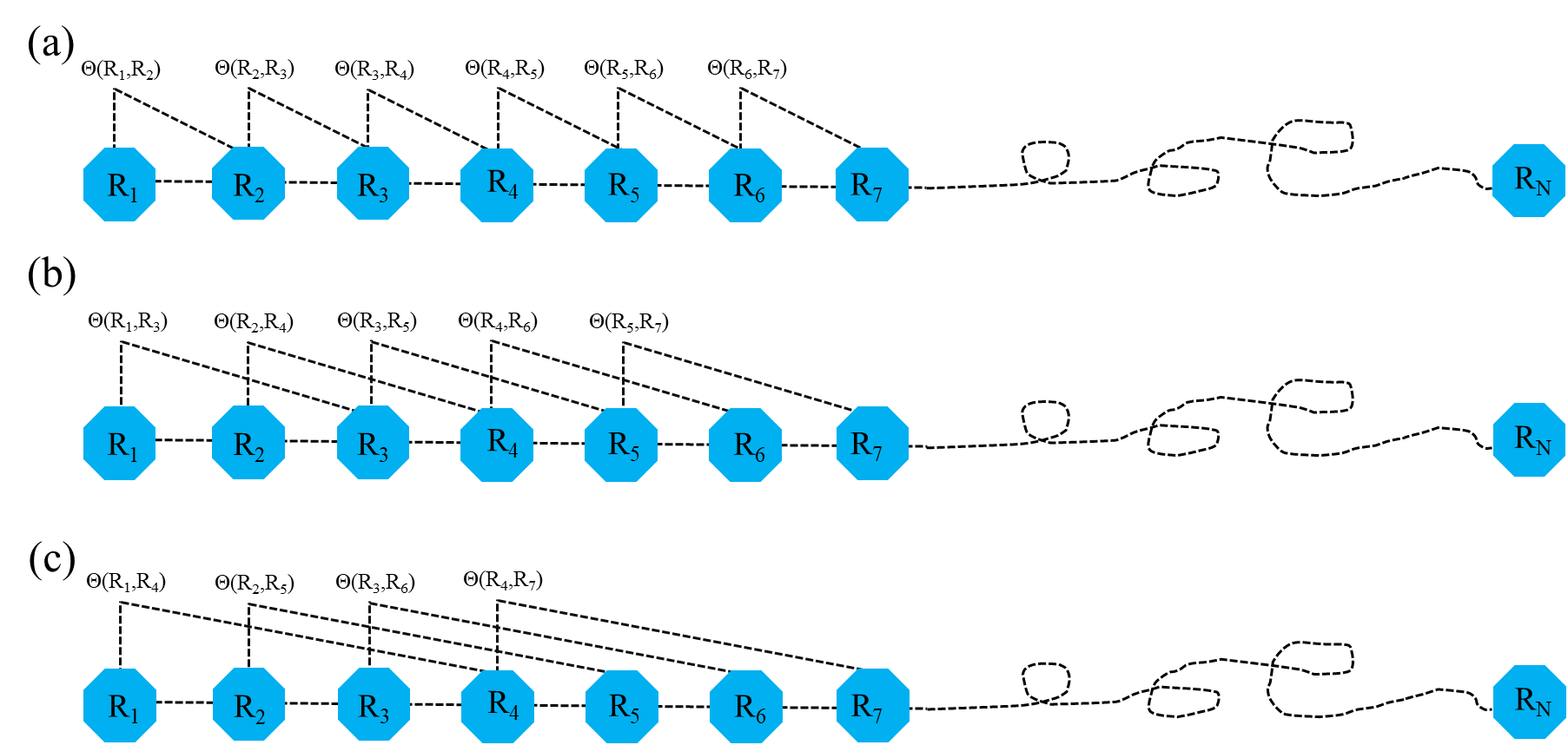

**Figure S8**. A schematic drawing to show (a) the first-tier, (b) the second-tier, and (3) the third-tier sequence order correlation mode along a protein sequence. (a) reflects the coupling mode between all the most adjacent residues, (b) shows the coupling between the adjacent plus one residues, and (c) shows the coupling between the adjacent plus two residues. This figure is adapted from (Chou, 2001) for illustration purposes.

For a set of *n* amino acid properties, it can be defined as:

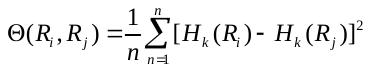

where *H_k_(R_i_)* is the *k*-th property in the amino acid property set for amino acid *R_i_*.

A set of descriptors called sequence order-correlated factors are defined as:

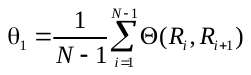

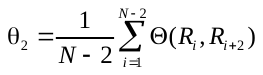

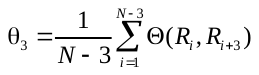

…

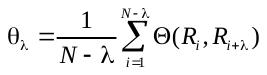

where λ (λ < *N*) is an integer parameter to be chosen. Let *f_i_* be the normalized occurrence frequency of amino acid *i* in the protein sequence. Then, a set of 20 + λ descriptors called the pseudo-amino acid composition for a protein sequence can be defines as:

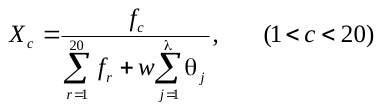

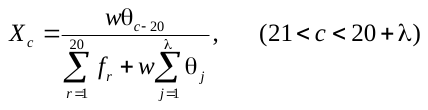

where *w* is the weighting factor for the sequence-order effect and is set to *w* = 0.05 in *iFeature* as suggested by Chou *et al*. (Chou, 2001).

### 1.15 PSSM profile (PSSM)

This feature descriptor (Cai, et al., 2012; Radivojac, et al., 2010) is extracted from the Position-Specific Scoring Matrix (PSSM) profile. An illustrated example of PSSM profile is provided in **Figure S10**. The PSSM profile can be obtained by running PSI-BLAST (Altschul, et al., 1997) against the uniref 50 database. The PSSM descriptor is usually applied to encode the peptides with equal length. Each amino acid in the peptide is represented by a 20-dimensional vector.

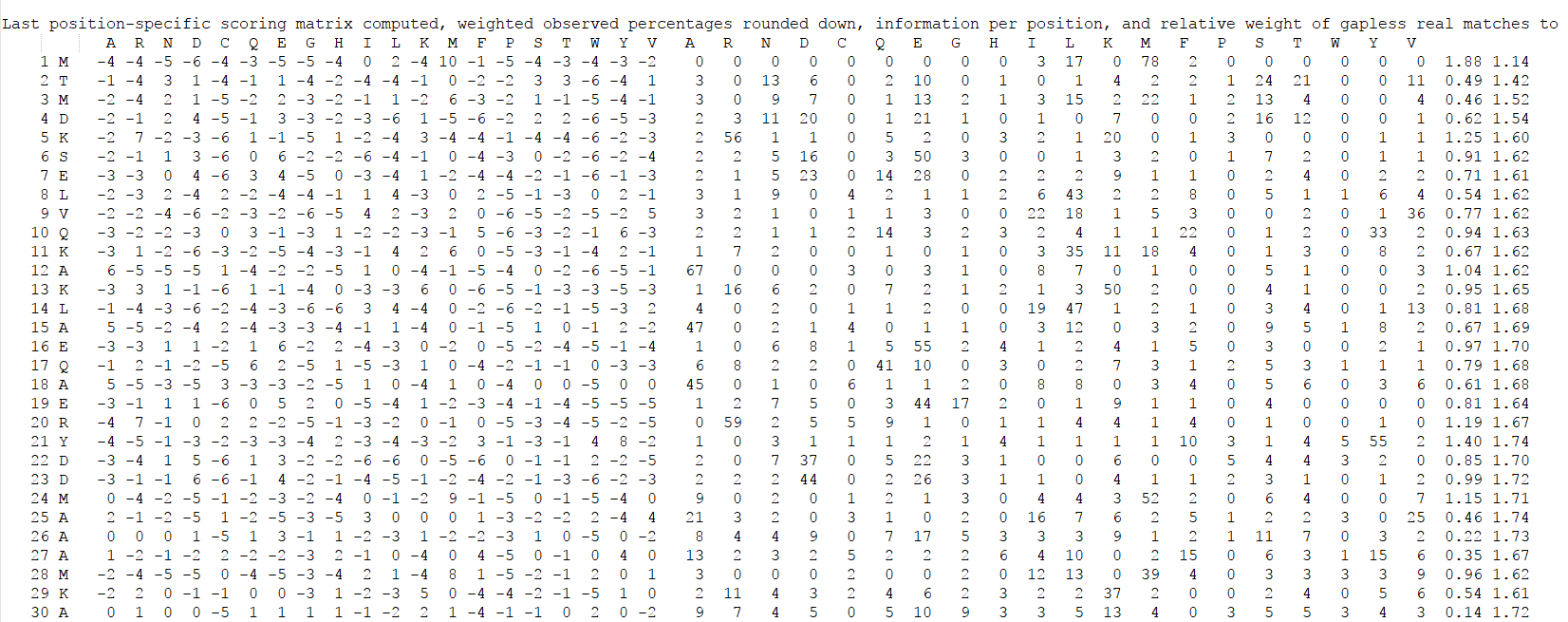

**Figure S10**. Example of a PSSM profile.

### 1.16 AAindex (AAINDEX)

Physicochemical properties of amino acids are the most intuitive features for representing biochemical reactions and have been extensively applied in bioinformatics research. The amino acid indices (AAindex) database (Kawashima, et al., 2008) collects many published indices representing physicochemical properties of amino acids. For each physicochemical property, there is a set of 20 numerical values for all amino acids. Currently, 544 physicochemical properties can be retrieved from the AAindex database. After removing physicochemical properties with value 'NA' for any of the amino acids, 531 physicochemical properties were left. In contrast to the residue-based encoding methods of amino acid identity and evolutionary information, a vector of 531 mean values is used to represent a sample for various window sizes. The AAINDEX descriptor (Tung and Ho, 2008) can be applied to encode peptides of equal length.
